# Interference with the HNF4-dependent gene regulatory network diminishes ER stress in hepatocytes

**DOI:** 10.1101/2023.02.09.527889

**Authors:** Anit Shah, Ian Huck, Kaylia Duncan, Erica R. Gansemer, Udayan Apte, Mark A. Stamnes, D. Thomas Rutkowski

## Abstract

In all eukaryotic cell types, the unfolded protein response (UPR) upregulates factors that promote protein folding and misfolded protein clearance to help alleviate endoplasmic reticulum (ER) stress. Yet ER stress in the liver is uniquely accompanied by the suppression of metabolic genes, the coordination and purpose of which is largely unknown. Here, we used unsupervised machine learning to identify a cluster of correlated genes that were profoundly suppressed by persistent ER stress in the liver. These genes, which encode diverse functions including metabolism, coagulation, drug detoxification, and bile synthesis, are likely targets of the master regulator of hepatocyte differentiation HNF4α. The response of these genes to ER stress was phenocopied by liver-specific deletion of HNF4α. Strikingly, while deletion of HNF4α exacerbated liver injury in response to an ER stress challenge, it also diminished UPR activation and partially preserved ER ultrastructure, suggesting attenuated ER stress. Conversely, pharmacological maintenance of hepatocyte identity *in vitro* enhanced sensitivity to stress. Several pathways potentially link HNF4α to ER stress sensitivity, including control of expression of the tunicamycin transporter MFSD2A; modulation of IRE1/XBP1 signaling; and regulation of Pyruvate Dehydrogenase. Together, these findings suggest that HNF4α activity is linked to hepatic ER homeostasis through multiple mechanisms.

## Introduction

Liver diseases kill approximately two million people worldwide every year, the majority of these from the sequelae of fatty liver disease (1). The major drivers of fatty liver disease are alcoholism, viral hepatitis, and, increasingly, obesity (2). It is now abundantly clear that ER stress is associated with fatty liver disease in humans and mouse models thereof (3). It is also likely that ER stress contributes to disease pathogenesis, since mice with an impaired ability to respond to ER stress in the liver are sensitized to experimental insults that lead to liver injury including alcohol, western diets, and pharmacotoxins that target the liver (4). The pathways by which ER stress affects hepatic function and exacerbates liver injury are still being characterized.

ER stress is caused by disruption to homeostasis in the organelle, most commonly in the form of impaired ER protein folding. Nascent secretory and membrane proteins are folded as they translocate into the ER lumen, and an imbalance between the load of such client proteins and the capacity of the organelle’s resident folding and modification machinery causes an accumulation of unfolded or misfolded proteins, leading to activation of the UPR. The UPR is a signaling cascade emanating from the ER, and its activation is diagnostic of ER stress. UPR signaling is attenuated when ER homeostasis improves, while it is largely perpetuated during severe ER stresses that cannot be overcome (5).

The essential function of the UPR is to restore ER homeostasis. The response accomplishes this directive by several mechanisms, the most widely conserved of which is the transcriptional upregulation of genes encoding ER chaperones and other factors that facilitate ER protein folding and trafficking (6). UPR signaling emanates from three ER-resident stress sensors—PERK (Protein kinase R-like ER Kinase), IRE1 (Inositol-Requiring Enzyme 1), and ATF6 (Activating Transcription Factor 6), each of which ultimately stimulates production of one or more transcription factors that activate expression of target genes. These transcription factors include ATF4 (for the PERK pathway), XBP1 (X Box Protein 1, for the IRE1 pathway), and ATF6 itself. The upregulation of ER chaperones occurs in every vertebrate organism and cell type in which the response has been examined. Yet the UPR also protects ER function by other means, including but not limited to: transient inhibition of protein synthesis though PERK-dependent phosphorylation of the translation initiation factor eIF2α (7); IRE1-dependent degradation of ER-associated mRNAs that encode ER client proteins (8); and stimulation of the autophagic turnover of the organelle itself (9). Thus, it is likely that additional pathways for protecting the ER emanate from UPR activation. Importantly, because the UPR is activated by ER perturbation and deactivated by the restoration of ER homeostasis, the response is likely “blind” to cellular conditions outside the organelle unless they impact ER function.

The best characterized transcription factors associated with the UPR are activators, and the vast majority of studies of the response have focused on the genes that are upregulated during ER stress. Conversely, despite the fact that transcriptional suppression is also observed by ER stress in every cell type, very little is known about the mechanisms by which this suppression is achieved and the purposes to which it is directed. We and others have observed that ER stress in the liver, commonly induced *in vivo* by the pharmacological agent tunicamycin (TM), leads to an extensive suppression of genes encoding metabolic regulators, and is accompanied by fat accumulation, or steatosis, in the liver. This steatosis is dramatically magnified when either UPR signaling or the ER protein folding capacity is impaired (10-13). Regulation of metabolic genes by the UPR suggests that there is an adaptive benefit to the ER in doing so. The canonical signaling molecules of the UPR and their downstream transcription factors—most of which are activators—are expressed in essentially all cell types that have been examined. Thus, this liver-specific regulation suggests that gene suppression is mediated not by direct repression by the transcription factors associated with the UPR, but instead by those factors interacting and interfering with liver-specific transcriptional machinery, as has been previously documented (14, 15).

One of the processes suppressed by the UPR in the liver is fatty acid oxidation, and we have shown that fatty acid oxidation stresses ER protein processing capacity by promoting hypooxidizing conditions within ER lumen. This effect occurs through the production of NADPH and the consequent reduction of glutathione that occur during TCA cycle flux (16, 17). Although this link between metabolism and ER homeostasis provides an example of how transcriptional suppression by the UPR might improve ER function, a bigger-picture view of what processes in the liver are suppressed by the UPR, and for what purpose, is largely unclear. Recently, it has been posited that the suppression of metabolic genes is part of a larger coordinated program by which the UPR suppresses the gene regulatory network (GRN) that controls hepatocyte identity (15).

Hepatocytes, which make up the large majority of the liver parenchyma, are considered “professional secretory cells” because of the abundant protein secretion required for their proper function, which is characterized by an extensive and elaborate ER network. Hepatocytes secrete albumin (the most abundant protein in the blood); lipoproteins including very low density lipoprotein (VLDL); antiproteases; coagulation factors; inflammatory mediators; and numerous other proteins (18). This extensive secretory need suggests that maintenance of hepatocyte identity is intrinsically tied to the functionality of the ER, and that ER stress could affect the hepatic differentiation state. Our previous work has suggested that many genes suppressed by ER stress in the liver might be targets of the transcription factor HNF4α (19). HNF4α is considered the “master regulator” of hepatocyte differentiation, and is essential for differentiation of hepatocytes (20, 21). It regulates numerous essential hepatocyte functions including metabolism, lipoprotein production, coagulation, and others (22). Loss of HNF4α causes hepatocytes to dedifferentiate and proliferate (23, 24), prevents the liver from functionally regenerating (25), and sensitizes the liver to diet-induced liver injury (26). Indeed, progressive loss of HNF4α function is a critical pathogenic component of a majority of liver diseases (27).

Building upon our previous work linking UPR activation to the HNF4α-dependent gene GRN (19), our goal here was to systematically examine the relationship between these two pathways, and to determine whether such a relationship conferred any functional benefit toward ER homeostasis. Here, we describe complementary bioinformatic, *in vivo*, and *in vitro* approaches that together demonstrate that ER stress impairs the HNF4α-dependent GRN, and that impairment of HNF4α in turn reduces the sensitivity of hepatocytes to ER stress.

## Results

### k-means clustering identifies a distinct group of genes involved in hepatocyte identify that is suppressed by persistent ER stress

We first sought to gain insight into the process of gene suppression by the UPR by testing whether these suppressed genes were coordinately regulated. To do this, we took advantage of published microarray data sets examining the global hepatic mRNA response to a tunicamycin (TM) challenge, in animals with a deletion of either ATF6α (10), IRE1α (13), or PERK (11). Although these experiments were performed by different groups at different times, they all used the same commercial microarray, the same dose and route of ER stress (1 mg/kg of TM injected intraperitoneally), and the same time-point (8 hours after injection). An additional data set, also using the same microarray, dose, and route of TM in wild-type or *Atf6α*-null animals but collected 34 hours after injection was also used (10). The premise of this approach was that genes whose regulation depended on a common factor would show a common pattern of regulation across these 4 experiments. Expression data for each probe set are provided in Table S1.

Among the genes *upregulated* by ER stress in the liver, the large majority depended on PERK signaling, while smaller sets of genes depended on ATF6 or IRE1, most of which were also PERK-dependent (Figure 1A), as previously reported in both the liver (11) and in fibroblasts (28). Among the genes *downregulated* by stress, approximately half could not be ascribed definitively to the action of one or more UPR pathways (Figure 1B). This result is likely a consequence of the fact that transcriptional upregulation occurs much more rapidly and robustly than transcriptional repression (29). Therefore, fewer genes show sufficiently robust differences in expression between wild-type and genetically-altered animals to pass the threshold of statistical significance. Notwithstanding that issue, the breakdown of genes whose suppression could be ascribed to one or more UPR pathways was nearly identical to that of the upregulated genes, with the large majority dependent on PERK (Figure 1B). Thus, these genes are likely suppressed by the known pathways of the UPR acting in unknown ways.

**Figure 1.**
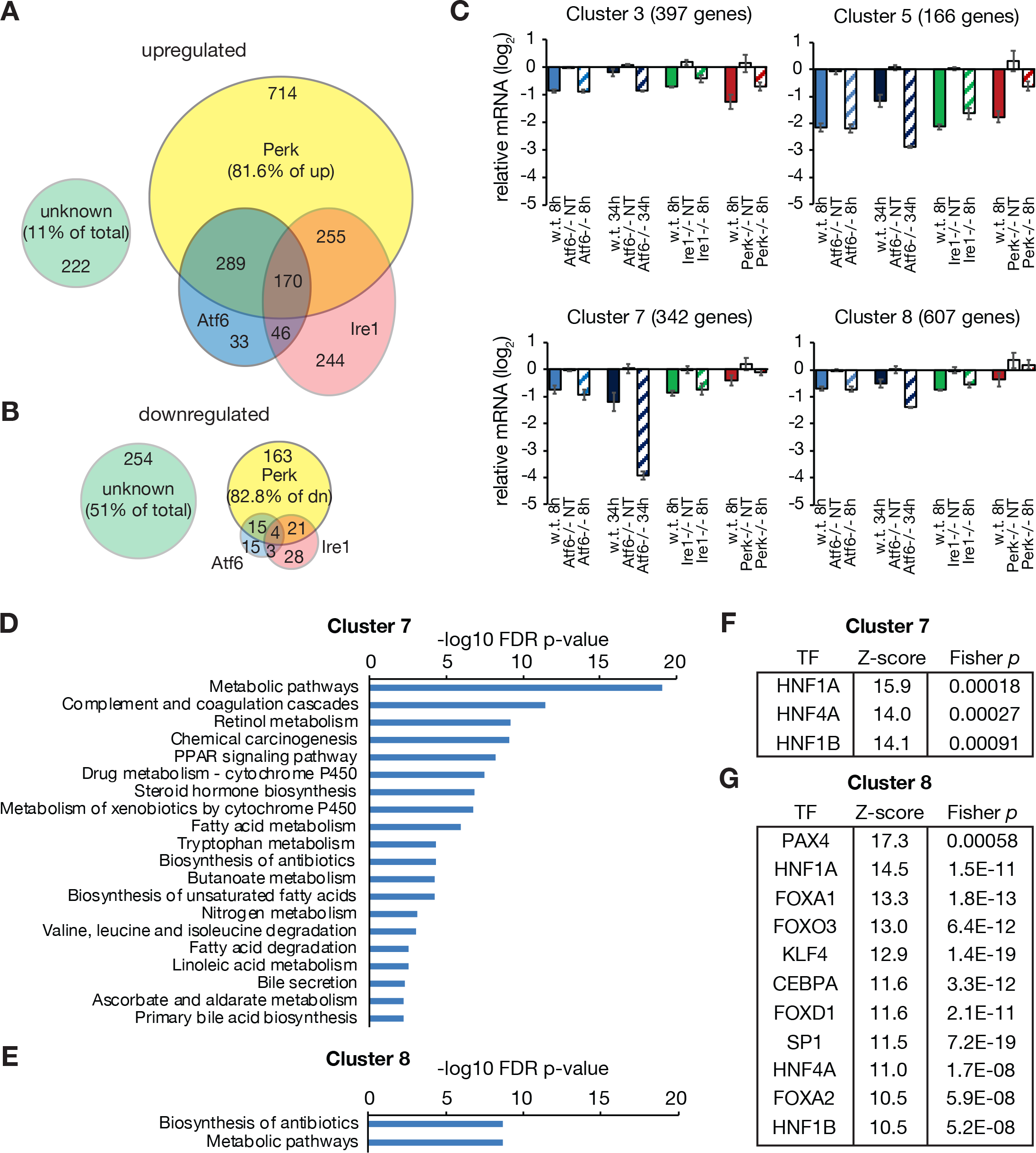
*k-means* clustering analysis of matched gene expression data sets from UPR mutants identifies a cluster of genes as likely to be HNF4α-dependent. **(A, B)** Data were assembled from published microarrays examining RNA expression in the livers of wild-type or *Perk*-, *Ire1α*-, or *Atf6α*-deleted animals following a challenge with 1 mg/kg TM. Individual genes were attributed to one or more pathways when loss of that pathway(s) abrogated the gene’s stress-dependent regulation. “Unknown” represents genes for which a clear assignment could not be made. Genes upregulated (A) and downregulated (B) by TM are shown. **(C)** *k-means* clustering was used to divide expression data from each microarray into 12 separate clusters based on the behavior of each gene in each group of each microarray. The centroids for each of the 4 clusters that captured downregulated genes are shown, along with the number of genes falling into each cluster. For each microarray, the expression for all groups was normalized against expression in wild-type animals treated with vehicle. The data are shown in log_2_-transformed format, so average expression in wild-type animals is “0”. As an illustration, the genes of Cluster 7 are suppressed, on average, ~16-fold by 34h TM treatment in *Atf6α*−/− livers. Upregulated cluster centroids are shown in Figure S1. **(D-G)** The genes of clusters 7 and 8 were subjected to pathway analysis using DAVID (D, E) and searched for enriched transcription factor binding sites (F, G) using oPPOSUM. Statistical cutoffs were FDR p < 0.01 for DAVID, and Z-score > 10 and Fisher score > 7 for oPPOSUM.

To identify potentially coordinately regulated genes among these datasets, we took an unbiased heuristic machine learning approach known as *k-means* clustering (30). This approach divides the full dataset into a prespecified arbitrary number of clusters, and then identifies centroids representing the average data values for each cluster within the dataset, seeking to identify the set of centroids that minimizes the average distance within each cluster. We tested 6, 12, 24, and 48 clusters, and found that 12 clusters provided enough flexibility for the algorithm to identify groups of genes with meaningfully distinct regulation patterns while mitigating the risk of overspecification. Among the 12 clusters (Table S2), upregulated genes comprised 8 (Figure S1) and downregulated genes comprised 4 (Figure 1C). These clusters— particularly the downregulated ones—were most notably distinguished by the magnitude of their regulation in wild-type animals (solid colored bars) and their responsiveness to longer-term ER stress in animals lacking ATF6α (black hashed bars). In particular, Cluster 7 stood out because the expression of those genes was profoundly suppressed (approximately 16-fold for the centroid) specifically by longer-term ER stress in *Atf6α*−/− animals (Figure 1C). This finding is notable because it is the context in which the extensive suppression of metabolic genes in the liver was first observed (10, 12).

Functional annotation of each of the 4 downregulated clusters supported the idea that Cluster 7 in particular was likely to be of particular interest. Cluster 7 was enriched for genes involved in a wide spectrum of biological processes that characterize functional hepatocytes, including nutrient metabolism, coagulation and complement cascades, drug metabolism, and bile production and secretion (Figure 1D). By contrast, only 2 pathways were identified for Cluster 8 genes—one of which was metabolism—and none of statistical significance was identified for Clusters 3 and 5 (Figure 1E and Table S3). In addition, the promoter regions (within 5kb of the transcriptional start site) of Cluster 7 genes were enriched for binding sites of the transcription factors HNF1α, HNF4α, and HNF1β (Figure 1F), which together drive hepatocyte differentiation—with HNF4α being higher in the gene regulatory network than HNF1α/β (31). Binding sites for these three transcription factors were also enriched among Cluster 8 genes, as were binding sites for other transcription factors implicated in hepatocyte differentiation and function including C/EBPα and members of the Forkhead family (Figure 1G) (32). In contrast, no enriched binding sites were found for Cluster 5, and Cluster 3 genes were not enriched for binding sites of transcription factors with obvious connections to metabolism or hepatocyte identity (Table S4). Taken together, these results suggest that there is a group of genes, comprising Cluster 7 and possibly 8, that are profoundly suppressed when ER stress cannot be remediated due to loss of ATF6. These genes participate in diverse hepatocyte functions beyond simply metabolism, and are likely direct or indirect HNF4α targets. This regulation is also mediated through PERK signaling, since only loss of PERK abrogated their suppression (Figure 1C).

### The transcriptional response to ER stress partially phenocopies loss of HNF4α

Since the genes of most interest in the above approach were putative HNF4α targets, we hypothesized that ER stress interferes with HNF4α activity. If this interference were the case, then we would expect that such HNF4α-dependent genes would be suppressed by either ER stress or loss of HNF4α, and, moreover, that ER stress would then have little or no further effect on those genes when HNF4α was absent. To test this idea, we injected *Hnf4α^fl/fl^* mice (25, 33) with AAV-TBG-GFP (*w.t*.), or AAV-TBG-CRE (34, 35) (*Hnf4α^LKO^*) to delete *Hnf4α* specifically in hepatocytes (Figure 2A), followed by injection with TM and sacrifice 14 hours thereafter. Global RNA expression was then profiled by RNA-seq (Table S5). Principal component analysis showed that both TM treatment and loss of HNF4α had substantial effects on global gene expression (Figure 2B). A heatmap of differentially regulated genes revealed several groups of interest (Figure 2C): Some genes (group (a)) were suppressed by ER stress independent of the presence or absence of HNF4α. Conversely, other genes (group (c)) were suppressed by loss of HNF4α independent of ER stress. However, a greater number of genes that were suppressed by loss of HNF4α were also suppressed by ER stress, but not then further suppressed by ER stress in the absence of HNF4α (group (b)). Interestingly, most of the genes upregulated by ER stress (group (d)) were upregulated to a lesser extent when HNF4α was lost. At least 20 percent of the genes significantly suppressed by ER stress were also significantly suppressed by loss of HNF4 and vice-versa (Figure 2D). That this overlap was not merely coincidental is attested by the fact that almost no overlap was observed among the genes upregulated by the two manipulations (see Figure 4A). Pathway analysis was used to identify the biological processes represented among genes downregulated by either ER stress or loss of HNF4α. This analysis revealed that, while the transcriptional response to loss of HNF4α was more functionally extensive than was the response to ER stress, all but one of the processes suppressed by ER stress was also suppressed by loss of HNF4α (Figure 2E)—and many of the genes involved the one process unique to ER stress, “liver development”, are nonetheless captured within the other processes regulated by loss of HNF4α.

**Figure 2.**
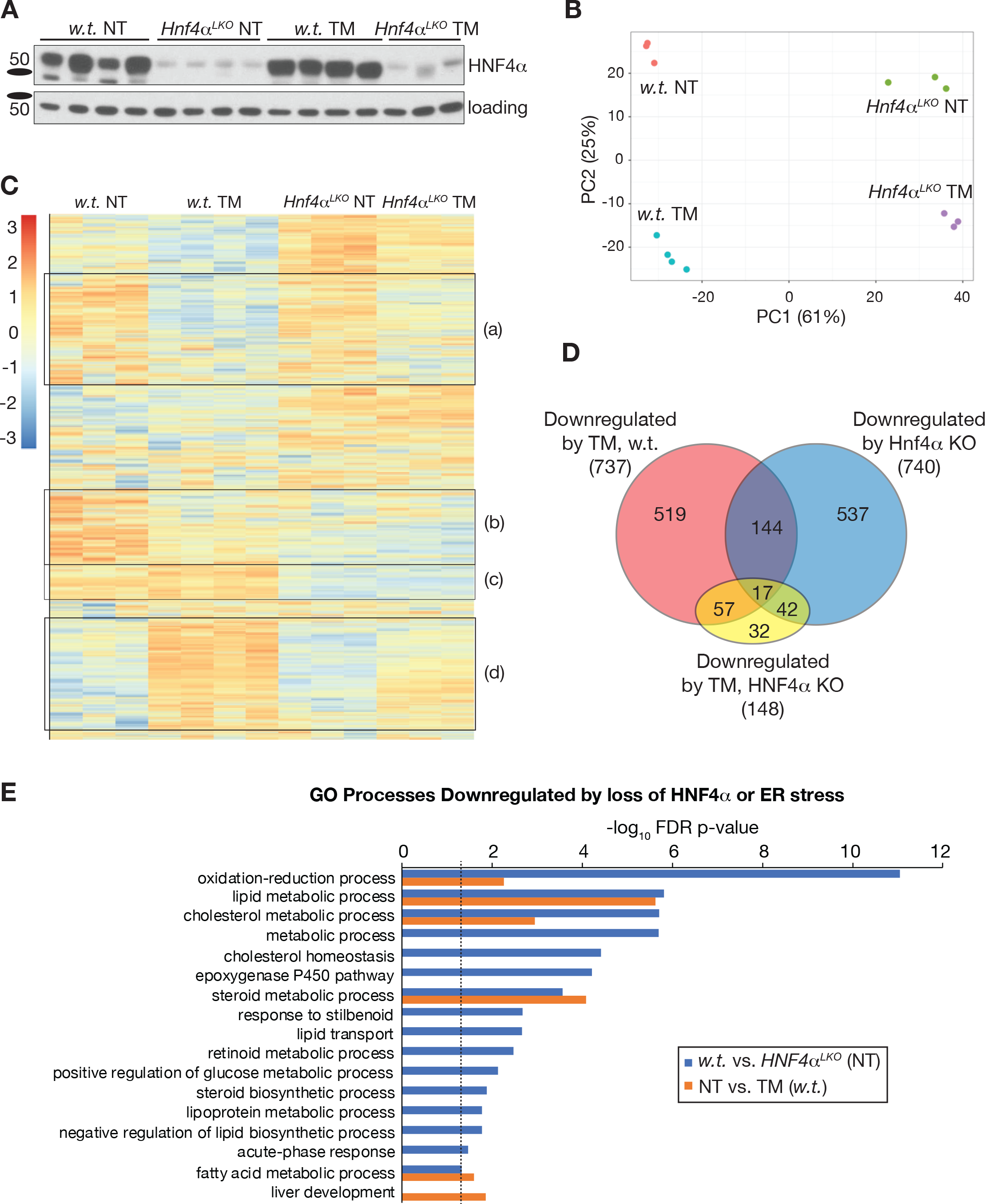
Liver-specific deletion of HNF4α partially phenocopies transcriptional suppression elicited by ER stress. **(A)** Deletion of HNF4α in the liver. Hnf4α^fl/fl^ mice were treated with AAV8-TBG-Cre (“*w.t.*”) or AAV8-TBG-eGFP (“*Hnf4α^LKO^*”) and challenged with 0.5 mg/kg TM, or vehicle control, for 14 hours as indicated. **(B)**Principal Component Analysis from RNA-seq. One sample each from the w.t. NT and NT *Hnf4α^LKO^* groups was excluded due to failure to pass pre-sequencing quality control. **(C)** Heatmap of differentially expressed genes in the liver. Groups of interest are noted (a-d) and described further in the text. **(D)** A Venn diagram shows a substantial overlap between genes downregulated by TM (red) or by loss of HNF4α (blue), and also shows that most of the genes downregulated by TM in wild-type animals are not further significantly downregulated when HNF4α is deleted (yellow). **(E)** Biological processes downregulated by the ER Stress or loss of *Hnf4α^LKO^* identified by gene ontology (GO) analysis.

Our *k-means* clustering identified a group of genes (Cluster 7) regulated by ER stress and presumed to be HNF4α targets, while the RNA-seq analysis identified genes that were suppressed separately by ER stress and loss of HNF4α (group (b)). Thus, we predicted that these two groups of genes would overlap substantially. Supporting this relationship, among the genes from Clusters 3, 5, 7, and 8 that were also identified as significantly suppressed by ER stress in the RNA-seq experiment, a higher percentage of those genes in Cluster 7 were also downregulated by loss of HNF4α compared to the other three clusters (41 out of 112, or 36.6 percent), and of these, the large majority (80.5 percent or 33 out of 41) were not further suppressed by ER stress when HNF4α was lost (Figures 3A and S2). Put in other terms, at least 29.4 percent (33 out of 112) of the genes from Cluster 7 belong to group (b) while clusters 3, 5, and 8 show less overlap (16.4, 22.2, and 9.1%, respectively). This finding reinforces the idea that Cluster 7 identifies HNF4α targets that are suppressed by ER stress.

**Figure 3.**
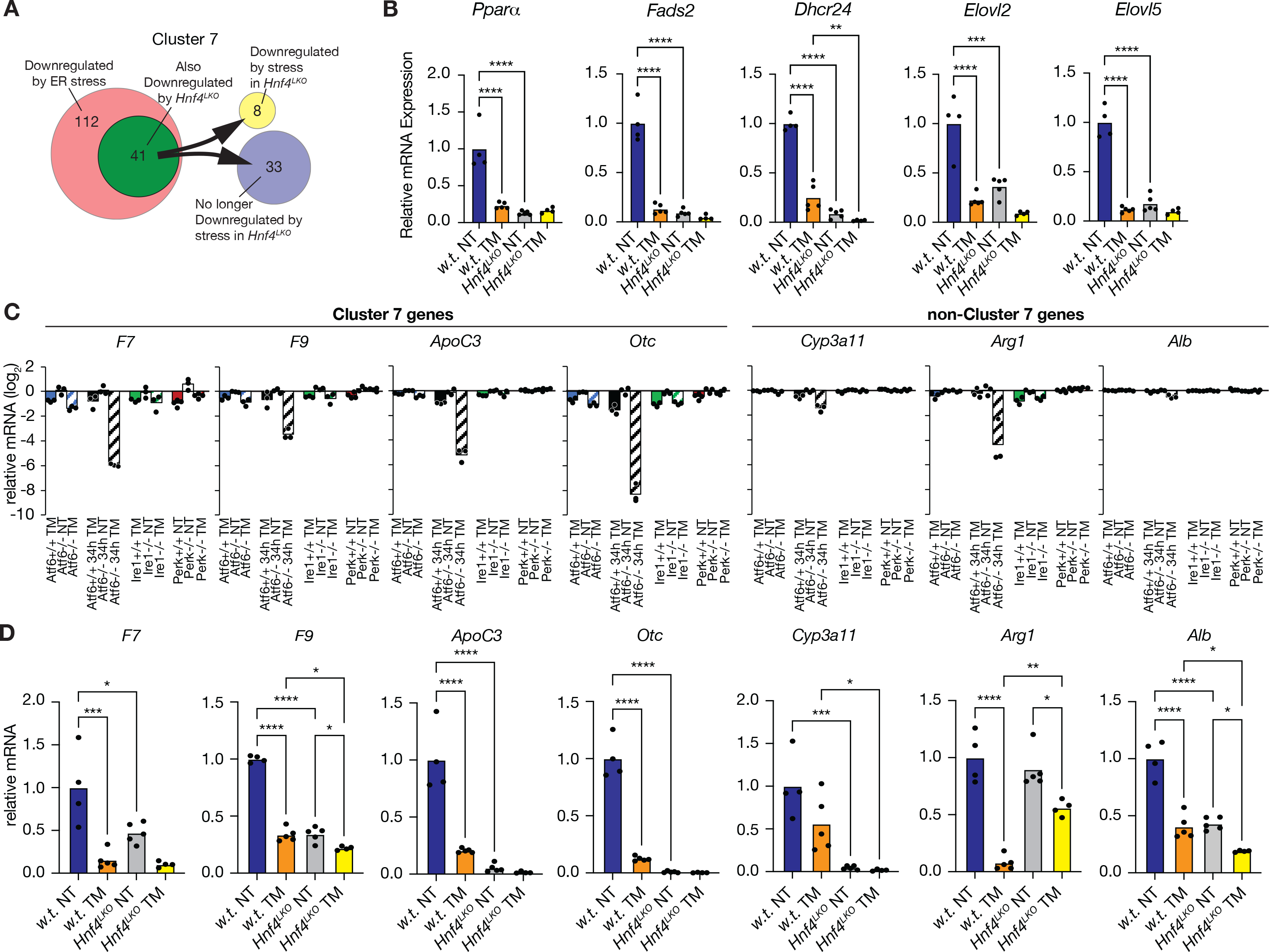
*Hnf4α*-dependent genes are selectively targeted by the ER stress. **(A)** Venn diagram illustrating that a substantial portion of Cluster 7 genes that were confirmed to be downregulated by TM in wild-type animals in the RNA-seq experiment were also downregulated by loss of HNF4α. Of these, the large majority (33/41) were not further downregulated by TM in *Hnf4α^LKO^* livers. Similar analysis for all four clusters of downregulated genes is shown in Figure S2. **(B)** Expression of representative Cluster 7 genes quantified by qRT-PCR from the livers of animals treated with TM similarly to those shown in Figure 2 (see Figure 4 and subsequent). Here and elsewhere, statistical comparison was by ANOVA with Tukey post-hoc comparison unless otherwise noted. *, *p* < 0.05; **, *p* < 0.01; **, *p* < 0.001; ***, *p* < 0.0001. **(C)** Expression of the indicated genes in individual samples from the microarray datasets shown in Figure 1. **(D)** qRT-PCR-determined expression of the genes in (C) determined as in panel (B).

To validate and extend these findings with maximal rigor, we next carried out an identical ER stress experiment, this time using both sexes of mice (only males were used for the RNA-seq experiment) and at a different location (in the lab of D.T.R., rather than the lab of U.A.). To examine gene regulation in this cohort with more granularity, we identified a group of 22 individual genes among the overlapping pathways identified in Figure 2E that were suppressed by both ER stress and loss of HNF4α. Of those 22 genes, 16 were also identified as downregulated in the initial *k-means* clustering analysis, and of those, 9 were in Cluster 7, 4 were in cluster 5, and 3 each were in Clusters 3 and 7 (Table S6; 3 genes were found in two clusters each based on different probe sets). We chose 5 of the Cluster 7 genes for further examination—*Pparα*, *Fads2*, *Dhcr24*, *Elovl2*, and *Elovl5*. The behavior of those genes in the microarray datasets is shown in Figure S3. Our prediction was that all 5 of these genes would be direct or indirect HNF4α targets, and that they would not be further significantly suppressed by ER stress when HNF4α was absent. Indeed, both predictions were true for all 5 genes (Figure 3B).

As a companion approach, we selected genes previously shown to be targets of HNF4α and to mark differentiated hepatocytes, including coagulation factors (*F7*, *F9*), urea cycle enzymes (*Arg1*, *Otc*), lipoprotein components (*ApoC3*), xenobiotic metabolism (*Cyp3a11*), and albumin (*Alb*) (21, 33, 36), without foreknowledge of how these genes would behave in either the *k-means* analysis or the targeted qRT-PCR analysis. Of these 7 genes, 4 were found within Cluster 7 (Figure 3C). From the RNA-seq analysis, each of these 4 was suppressed by both ER stress and by loss of HNF4α, and only 1—*F9*—was significantly further suppressed by ER stress in *Hnf4α^LKO^* animals, albeit to a very modest extent (Figure 3D). Among the genes not identified as being in Cluster 7, *Cyp3a11* was not significantly suppressed by ER stress and *Arg1* was not significantly suppressed by loss of HNF4α, while *Alb* behaved similarly to *F9*. Together, the data in Figures 1-3 suggest that, while Cluster 7 does not identify all HNF4α targets, and not all HNF4α targets are suppressed by ER stress, the genes of Cluster 7 are strongly enriched for a group of HNF4α targets that contribute to hepatocyte identity and are coordinately suppressed by ER stress in an HNF4α-dependent manner. This group emerges despite the inherent noisiness attendant in grouping genes from whole transcriptome datasets.

### Loss of HNF4α diminishes ER stress sensitivity

Even though loss of HNF4α is deleterious to the liver, the observation that its activity is seemingly impaired by the UPR implies that there is a functional benefit to the hepatocyte ER from this impairment during ER stress. The first observation in support of this idea was the effect of HNF4α disruption on *bona fide* UPR targets—those genes that are robustly upregulated by ER stress in wild-type animals. Notably, even though very few of these genes are responsive to loss of HNF4α alone, the majority of them (58.6 percent or 519 out of 885) were no longer significantly <u>upregulated</u> by ER stress in *Hnf4α^LKO^* livers in our RNA-seq experiment (Figure 4A). This finding was supported by Gene Set Enrichment Analysis (GSEA); the Unfolded Protein Response gene set was significantly suppressed in TM-treated *Hnf4α^LKO^* livers compared to wild-type TM-treated livers (Figure 4B) meaning that, collectively, these genes were expressed to a lower extent in the latter group than in the former one. Illustrating this point more precisely, of the 20 genes most strongly upregulated by ER stress in wild-type livers, a nearly uniform diminishment of that upregulation was seen in *Hnf4α^LKO^* livers (Figure 4C). These findings were supported by our companion experiment in the second cohort of animals (Figure 4D) where, qualitatively, there was lower upregulation in *Hnf4α^LKO^* livers of UPR target genes including *Bip*, *Edem1*, *Derl3*, *Erp72*, and others (Figure 4E and data not shown). This finding supports the conclusion that UPR signaling is diminished in the livers of animals lacking HNF4α. Yet, interestingly and in contrast, the splicing of *Xbp1* mRNA, which occurs as a consequence of IRE1 activation, was enhanced by loss of HNF4α (Figure 4F).

**Figure 4.**
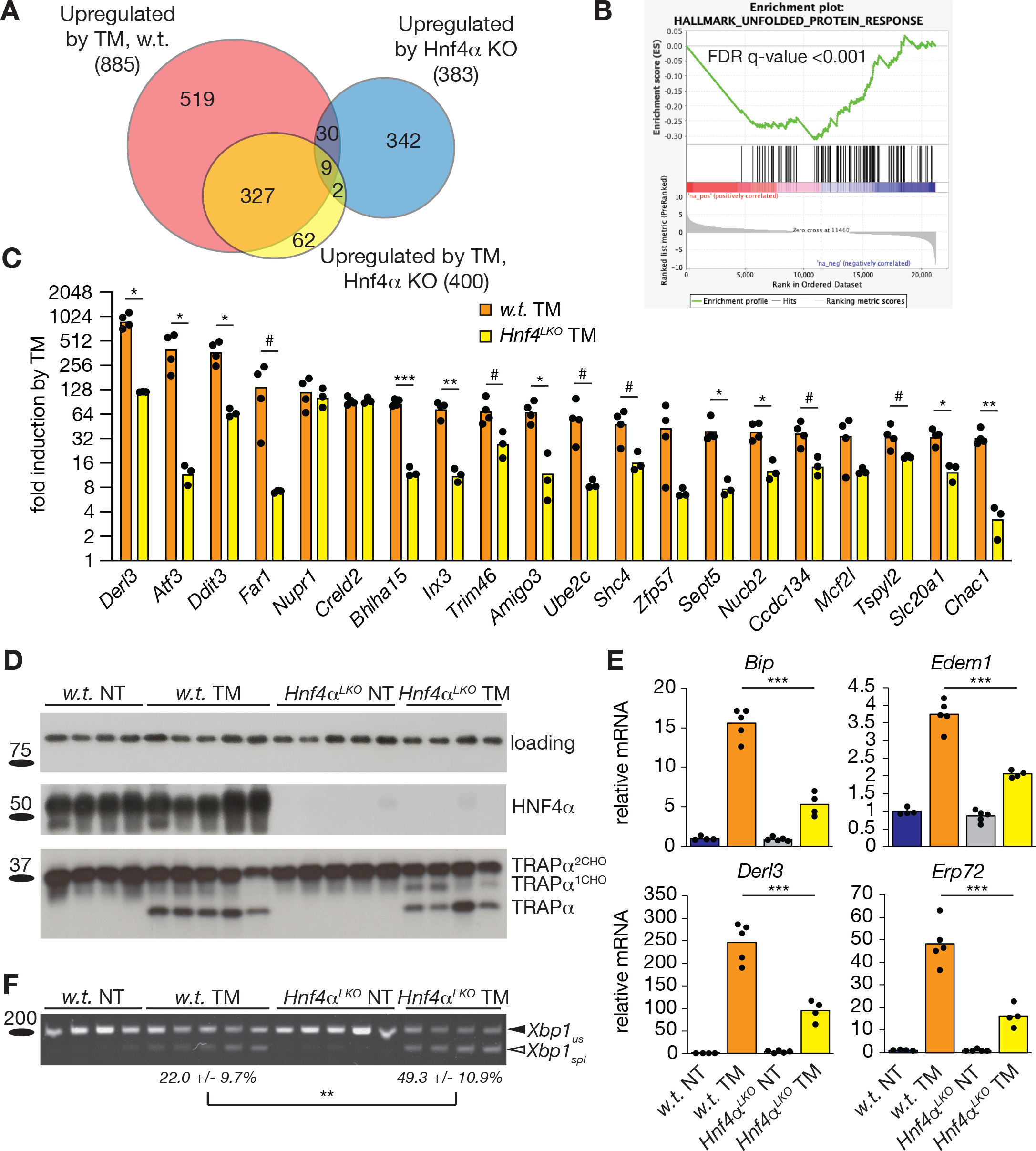
Deletion of HNF4α in the liver diminishes UPR sensitivity during ER stress. **(A)** Analysis of genes upregulated by TM from RNA-seq data, analyzed similarly to Figure 2D. **(B)** Gene Set Enrichment Analysis (GSEA) for Unfolded Protein Response genes in TM-treated *Hnf4α^LKO^* animals compared to TM-treated wild-type animals, showing consistently lower expression in the former group. **(C)** Fold-induction by TM in both genotypes of the 25 genes most highly upregulated in wild-type livers. Statistical comparison was by one-way ANOVA followed by Benjamini-Hochberg adjustment for multiple comparisons. #, *p* between 0.05 and 0.1. **(D)** Immunoblot showing effective deletion of HNF4α and inhibition by TM of glycosylation of the ER-resident glycoprotein TRAPα, in an experiment conducted similarly to that shown in Figure 2. **(E)** qRT-PCR analysis of the indicated UPR target genes from the experiment shown in (D) **(F)** *Xbp1* mRNA splicing, assessed by conventional RT-PCR with the spliced (“spl”) and unspliced (“us”) forms indicated

Diminished UPR signaling could be accounted for either by improved ER homeostasis in *Hnf4α^LKO^* livers or by neutered UPR signaling irrespective of ER homeostasis. In fact, neutered UPR signaling would be likely to exacerbate ER disruption caused by a stressor due to the lack of an adequate protective response. To discriminate between these possibilities, we examined ER ultrastructure by transmission electron microscopy. Loss of HNF4α on its own had no discernible effect on ER ultrastructure. Conversely, TM caused dramatic disruption of lamellar ER morphology in both genotypes. In wild-type animals, the ER completely vesiculated. However, this disruption was less severe in *Hnf4α^LKO^* livers, as the ER did not vesiculate as completely as in wild-type animals, and retained some partially lamellar organization, particularly in areas adjacent to mitochondria (Figure 5A). To measure this change, ER circularity was quantified from the experiment shown in Figure 5A, with the measurer blinded to sample identity; this analysis confirmed a less extensive circularization— i.e., vesiculation—of the ER in *Hnf4α^LKO^* livers (Figure 5B). Also noted was a slight but significant elongation of mitochondria that was present upon ER stress only in *Hnf4α^LKO^* livers (Figure 5C), which suggests that loss of HNF4α makes mitochondria responsive to ER stress in a way that they are not in wild-type animals.

**Figure 5.**
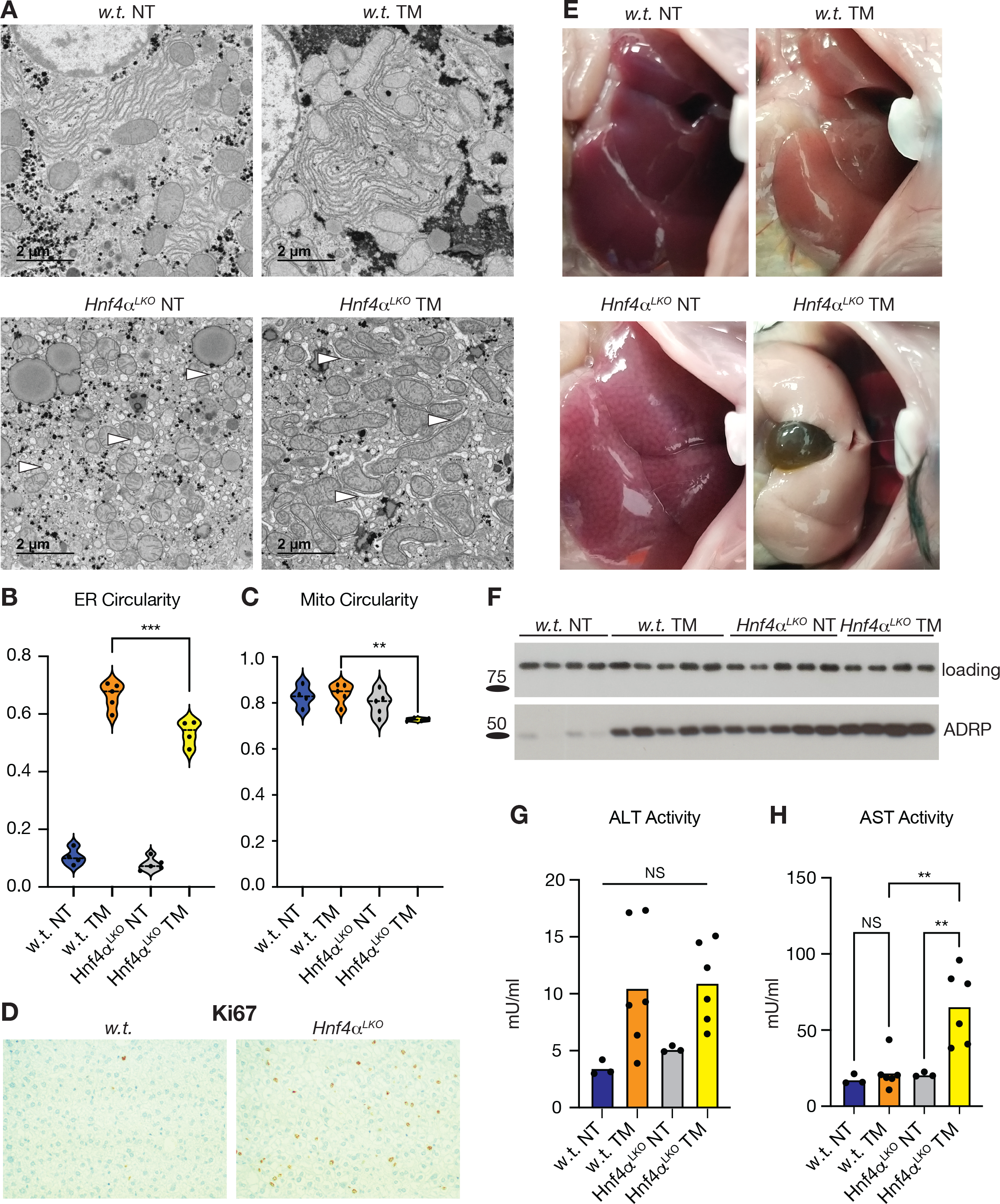
Loss of HNF4α aggravates liver injury while protecting the ER. **(A)** Representative TEM images showing ER and mitochondrial morphology in w.t. and *Hnf4α^LKO^* liver treated with vehicle or TM as in Figure 4. Arrowheads highlight ER structures in TM-treated samples. **(B, C)** ER (B) and mitochondrial circularity (C) were quantified from EM images with the quantifier blinded to image identity. The degree of circularity was averaged over multiple images from each sample, which those averages shown as points within the violin plots. Statistical comparison was carried out on a per-sample basis, not a per-image basis. **(D)** Ki67 immunostaining shows hepatocellular proliferation in *Hnf4α^LKO^* livers. **(E)** Profound steatosis and cholestasis are observed specifically in *Hnf4α^LKO^* livers after a 48h TM challenge. **(F)** The lipid droplet marker protein ADRP is elevated by both TM (14h) and HNF4α deletion, and further elevated by their combination. **(G, H)** Plasma AST (G) and ALT (H) in w.t. and *Hnf4α^LKO^* animals treated with vehicle or TM for 48h.

Despite apparently protecting ER homeostasis, loss of HNF4α exacerbated signs of liver injury upon ER stress challenge. As expected (21), loss of HNF4α led to hepatocellular proliferation, seen by Ki67 staining (Figure 5D), that was independent of ER stress treatment (data not shown). In addition, while both ER stress and loss of HNF4α led to hepatic lipid accumulation, the combined effect of both manipulations was, after a longer (48 hour) challenge, a profound hepatic steatosis and an enlarged and discolored gall bladder that was much more prominent in *Hnf4α^LKO^* TM-challenged animals than wild-type (Figure 5E). Additional evidence for hepatic steatosis was seen in expression of the lipid droplet marker protein ADRP, which was increased by both TM and loss of HNF4α and increased further by the combination of these manipulations (Figure 5F). While loss of HNF4α did not affect plasma ALT (Figure 5G), AST was elevated only in *Hnf4α^LKO^* animals treated with TM (Figure 5H). Together, the data in Figure 5 suggest that loss of HNF4α protects the ER organelle from TM challenge at the expense of the organ.

### Maintenance of hepatocyte specification sensitizes cells to ER stress in vitro

Primary hepatocytes begin dedifferentiating immediately upon isolation and culture, as seen by their change to a more fibroblastic morphology (Figure 6A) and loss of HNF4α expression (Figure 6B). This propensity proved to be extremely vexing to our attempts to recapitulate *in vitro* the effects of *in vivo* deletion of HNF4α. The response of primary hepatocytes to ER stressors and to overexpression or knockdown of HNF4α was highly variable from experiment to experiment, which is perhaps unsurprising given how markedly HNF4α expression is lost in these cells after only one day in culture, not to mention the many other cellular changes that likely accompany this dedifferentiation.

**Figure 6.**
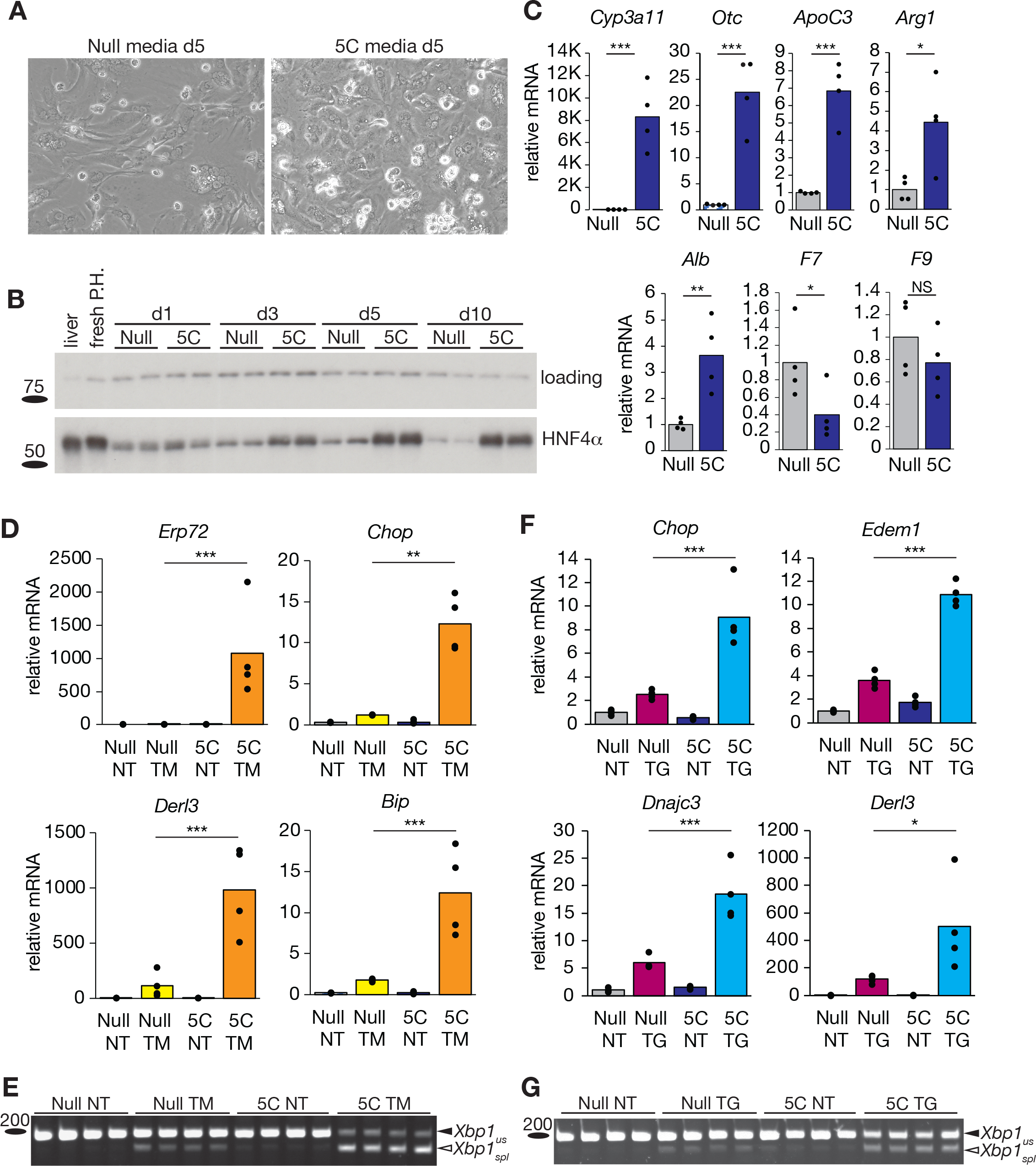
Pro-differentiation 5C media sensitizes primary hepatocytes to ER stress. **(A)** Brightfield images of hepatocytes cultured for 5 days in either null or 5C-containing media, with the null cells showing a more fibroblastic appearance and the 5C cells retaining a more cobblestone-like morphology. **(B)** Immunoblot showing reversal of HNF4α suppression in hepatocytes cultured in 5C media. **(C)** qRT-PCR showing expression of a set of genes diagnostic of hepatocyte differentiation in null-compared to 5C-containing media. **(D, E)** 5C-cultured cells are more sensitive to TM, as shown by qRT-PCR analysis of UPR target genes (D) and Xbp1 mRNA splicing (E). **(F, G)** Same as (D, E) except using thapsigargin (TG) as the ER stressor rather than TM.

However, it was recently demonstrated that a cocktail of 5 compounds (5C media) containing agonists and antagonists of various signaling pathways important in the maintenance of hepatocyte identity could forestall the dedifferentiation of primary human hepatocytes *in vitro* (37). This 5C cocktail was partially effective in mouse primary hepatocytes as well, preserving the more cobblestone-like morphology of hepatocytes and reversing the decline in HNF4α expression seen upon culture (Figure 6A, B). The 5C-containing media also enhanced the expression of most (but not all) of the hepatocyte identity genes examined in Figure 3 (Figure 6C)—albeit to levels in most cases still well below those seen in the liver (data not shown).

We reasoned that this method could be used to leverage the natural dedifferentiation of hepatocytes in culture into an independent test of whether the differentiation state of hepatocytes affected cells’ sensitivity to stress. Indeed, cells grown in 5C-containing media were more sensitive to TM, as seen in upregulation of UPR target genes including *Erp72*, *Chop*, *Derl3*, and *Bip* (Figure 6D). In contrast to the HNF4α knockout *in vivo*, in these cells, splicing of *Xbp1* mRNA followed the same pattern as UPR gene expression, in that null-treated cells showed less splicing (Figure 6E). Similar results were obtained with the independent ER stressor thapsigargin (TG) (Figure 6F, G). These results are consistent with the idea that hepatocytes in which HNF4α activity and hepatocyte identity are suppressed are less sensitive to ER stress.

### Multiple pathways link HNF4α to ER stress sensitivity

We next explored potential mechanisms linking HNF4α to ER homeostasis. One possibility was that loss of HNF4α interfered with the pharmacological potency of TM. Our readout for the efficacy of TM was inhibition of glycosylation of the ER-resident glycoprotein TRAPα. TRAPα has two N-linked glycosylation sites, and while essentially all of the hypo-glycosylated TRAPα in wild-type animals was completely devoid of N-linked sugars, a noticeable fraction of the TRAPα in *Hnf4α^LKO^* animals was instead mono-glycosylated (Figure 4D). This observation raised the possibility that TM was less potent in *Hnf4α^LKO^* animals than in wild-type. Indeed, deletion of HNF4α profoundly suppressed expression of mRNA encoding the TM transporter MFSD2A (Figure 7A) (38). However, while *Mfsd2a* expression was elevated in TM-treated cells grown in 5C media compared to cells grown in null media, the 5C treatment suppressed basal expression of the gene (Figure 7B). In addition, apparent changes in TRAPα glycosylation might be due not to differences in the pharmacokinetics of TM but in changes to the synthesis and turnover of TRAPα itself, since TM only affects the glycosylation of nascent glycoproteins. Therefore, while suppression of *Mfsd2a* expression possibly contributes to the diminished stress observed upon TM treatment *in vivo*, other pathways need to be invoked *in vitro* and possibly *in vivo* as well.

**Figure 7.**
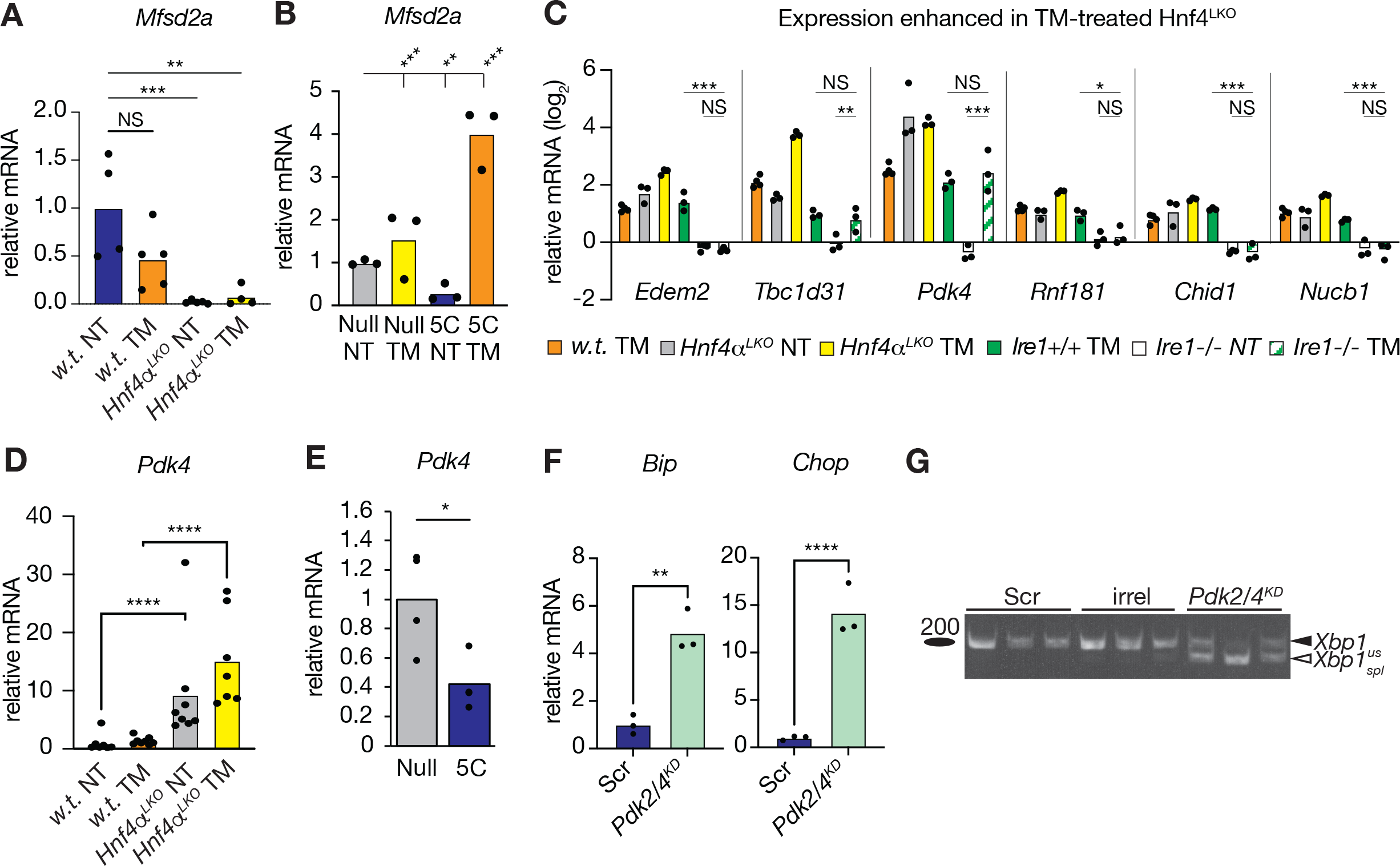
Multiple potential pathways link HNF4α to ER homeostasis. **(A, B)** qRT-PCR showing expression of *Mfsd2a* in (A) wild-type and *Hnf4α^LKO^* livers and (B) null- or 5C-cultured hepatocytes, treated with vehicle or TM. **(C)** Aggregated expression data from RNA-seq (orange, gray, and yellow bars) and microarray (green, white, and green-hashed bars) of genes whose expression was most elevated in TM-treated *Hnf4α^LKO^* TM-treated livers relative to w.t. TM-treated livers. As in Figure 1, expression is normalized against w.t. untreated livers, and is shown on a log_2_ scale. **(D, E)** Expression of *Pdk4* in (D) wild-type and *Hnf4α^LKO^* livers and (E) null- or 5C-cultured hepatocytes treated with vehicle or TM. In (D), data were combined from both *in vivo* experiments. **(F, G)** Expression of the UPR target genes *Bip* and *Chop* (F) and *Xbp1* mRNA splicing (G) in primary hepatocytes in which *Pdk2* and *Pdk4* are knocked down.

Another likely contributing pathway is the apparent link between HNF4α and IRE1 activation, as seen in *Hnf4α^LKO^* animals *in vivo* (Figure 4F). The IRE1/XBP1 axis is broadly protective against ER stress in general and more specifically in the liver (3). While the expression of most TM-induced genes was attenuated in TM-treated *Hnf4α^LKO^* animals (Figures 2C and 4A), this was not universally true. Among the genes showing the most significantly *elevated* expression in TM-treated *Hnf4α^LKO^* animals compared to TM-treated wild-type animals, several (*Edem2*, *Rnf181*, *Chid1*, and *Nucb1*) were fully dependent upon IRE1 activity for their induction (Figure 7C). Notably, all of these genes were also elevated solely by deletion of HNF4α (compare gray bars to the baseline), and even further upregulated in TM-treated *Hnf4α^LKO^* animals (compare yellow bars to orange bars), which is consistent with the idea that IRE1/XBP1 signaling is elevated both basally and in response to ER stress in *Hnf4α^LKO^* animals.

One of the few IRE1-*independent* genes that was strongly induced by HNF4α deletion was *Pdk4* (Figure 7D), which encodes an isoform of Pyruvate Dehydrogenase Kinase. This enzyme deactivates Pyruvate Dehydrogenase, thus inhibiting conversion of pyruvate into acetyl-CoA and, consequently, TCA cycle flux (39). Mirroring this observation, 5C treatment decreased expression of *Pdk4* (Figure 7E). The other liver-expressed isoform of PDK, *Pdk2*, was unaffected in both conditions (Figure S4A, B). We have previously demonstrated that attenuation of TCA cycle flux protects hepatocytes from ER stress (16). Consistent with that idea, combined knockdown of *Pdk2/4* (Figure S4C) was sufficient to cause ER stress, as seen by upregulation of UPR target genes (Figure 7F, S4D) and splicing of *Xbp1* (Figure 7G). Thus, the inverse association between HNF4α and PDK4 might provide an additional mechanism by which loss of HNF4α protects against ER stress.

## Discussion

It has long been understood that the major function of the UPR is the restoration of ER homeostasis, which it effects primarily through transcriptional regulation. Across eukaryotic organisms and cell types, the core of this response is the upregulation of ER chaperones and other factors that facilitate ER protein folding, trafficking, and degradation (40, 41). Yet the transcriptionally suppressive facet of the UPR is much less understood, and much less conserved from one cell type to another. In the liver, this response includes the suppression not only of genes involved in metabolism, as we and others have shown, but also more broadly of genes that demarcate hepatocyte identity as shown more recently (15) and here by us.

The principal goal of this study was to understand the nature and consequences of this suppressive program for the hepatocyte. Our major conclusions are that ER stress interferes with a substantial portion of the HNF4α-dependent GRN, and that this interference likely confers to the cell a degree of resistance to ER stress even at the potential cost of the hepatocyte functions to which HNF4α activity is crucial. Although HNF4α has previously been linked to the ER stress response by us and others (15, 19), this work is the first direct demonstration that HNF4α affects ER stress sensitivity *in vivo*. That the interference of hepatocyte identity protects the ER was seen using two complementary models—one based on *in vivo* deletion of HNF4, and one based on the pharmacological forestalling of dedifferentiation *in vitro*. In both cases, diminished ER stress signaling was seen when HNF4α was absent or attenuated.

Although the responsiveness of numerous HNF4α-dependent genes to ER stress suggests that there is a discrete pathway by which the UPR interferes with HNF4α activity, the nature of this pathway is not yet clear. The simplest possibility, suggested elsewhere (15), is if HNF4α is itself a target of UPR-directed transcriptional suppression. However, in both this paper (Figs. 2A, 4D) and our previous work (19), ER stress-induced suppression of HNF4α targets preceded any detectable changes in HNF4α protein expression. Therefore, it seems more likely that the UPR interferes with HNF4α activity, and that effects on HNF4α expression are a consequence of the autoregulatory feedback loop by which HNF4α regulates its own expression (42). The major remaining possibilities are that the UPR prevents HNF4α from binding to its target promoter/enhancers or that it impairs its activity when bound. The observation that most but not all of the genes suppressed by loss of HNF4α are also suppressed by ER stress (groups (b) vs. (c), Fig. 2C) suggests that HNF4α retains its activity toward some of its target genes, and thus that wholesale loss of its DNA binding capacity during ER stress is unlikely. For this reason, we favor, on principle, models whereby the UPR interferes with HNF4α activity even when it is capable of binding its target sites. This question can ultimately be resolved by examining global HNF4α chromatin binding and its interactions with the transcriptional machinery during stress.

How might the UPR interfere with HNF4α in ways besides preventing its DNA binding? The fact that the suppression of genes of Cluster 7 is—like most of the other stress-suppressed genes—PERK-dependent, points to a role for either translational repression mediated by eIF2α phosphorylation or to one or more of the factors that is translationally upregulated by eIF2α phosphorylation such as ATF4, CHOP (C/EBP Homologous Protein), or others (43, 44) (or possibly to both mechanisms). As we have shown, the stress-dependent repression of the Cluster 7 gene *Ppara* is lost when CHOP is deleted (45). As an inhibitory C/EBP-family transcription factor, CHOP could act by dimerizing with another C/EBP-family member that would ordinarily costimulate gene expression in concert with HNF4α, and squelching it (46). An attractive candidate would be C/EBPα, which is known to coregulate many genes with HNF4α (47). However, we found that liver-specific deletion of C/EBPα alone had only modest effects on the expression of Cluster 7 genes (data not shown). An alternate possibility is that CHOP binds to C/EBP binding sites (as we have shown (45)) and exerts a dominant negative influence on the transcription of genes that are ordinarily coregulated by C/EBPα and HNF4α. Whether there is a wider role for CHOP in linking ER stress to HNF4α activity, and whether such an effect does or does not depend on loss of C/EBPα, remain under investigation. We also note that ER stress suppresses genes that are not HNF4α targets (group (a), Fig. 2C), showing, unsurprisingly, that there are additional stress-dependent mechanisms for repressing hepatic gene expression as suggested (15).

Our findings show that loss of HNF4α is beneficial to ER functionality at least during a TM challenge. They suggest that, more broadly, partially unwiring the hepatocyte gene expression program might be exercised by the UPR in order to protect ER function during physiological challenges. It stands to reason that the ER in hepatocytes—with their need for abundant protein secretion—might be intrinsically more taxed than in cells in which the HNF4α-dependent program has been (temporarily) silenced. Indeed, the fact that liver development requires an intact UPR (48) suggests that the differentiation of hepatocytes, like other so-called “professional secretory cells”, places a stress on the ER due to the secretory protein burden. It might be that interruption of HNF4α activity simply diminishes production of secretory pathway client proteins (albumin, lipoproteins like APOC3, clotting factors like F7 and F9, etc.), thereby reducing the burden on the hepatocyte ER. We found, however, that cells cultured in null media, though they were relatively resistant to ER stress (Figure 6), did not in fact have a reduced total load of nascent ER client proteins compared to 5C-treated cells (data not shown). This finding suggests that, at least in that model system, some other mechanism must drive the increased stress sensitivity of 5C-treated cells.

A completely different mechanism by which loss of HNF4α might protect ER function is through control of the TM transporter MSFD2A. Testing this idea was challenging because hepatocytes are largely refractory to other common ER stressors like TG or dithiothreitol (as can be seen from the very modest response of null-cultured cells to a high dose of TG in Figure 6F), making differences in sensitivity difficult to detect. It is surprising, given how strongly *Msfd2a* is suppressed by HNF4α deletion *in vivo* (Figure 7A), that there appears to be very little if any effect on the actions of the drug at least on the glycosylation of the ER-resident glycoprotein TRAPα, for which the difference between genotypes was minor at most (Fig. 4D). These results hint either that MSFD2A is expressed at levels far above what is necessary to transport TM (obviously not the physiological substrate of the channel), or that another means of TM transport in the liver exists—or both.

It is perhaps more likely that an interaction (in the broad, genetic sense) between HNF4α and IRE1 signaling confers at least some of the resistance to ER stress when HNF4α is absent. Splicing of *Xbp1* was elevated in *Hnf4α^LKO^* livers (Fig. 4F), in marked contrast to other readouts of UPR output, which were diminished. Our findings are, to our knowledge, the first demonstration of a functional relationship between IRE1 and HNF4α. While the nature of this relationship is still being investigated, we did find that the gene encoding HSP70 (*Hsp1a1*) was upregulated specifically in *Hnf4α^LKO^* livers (data not shown). It has previously been shown that HSP70 can bind to the cytosolic domain of IRE1 and enhance its activity towards *Xbp1* (49). However, no apparent unique linkage between HNF4α expression and *Xbp1* splicing was observed in the 5C system.

A feature common to both systems was the upregulation of *Pdk4* in cells lacking HNF4α (Fig. 7D, E). Having previously demonstrated that pharmacological inhibition of PDKs causes ER stress (16), and here showing the same with knockdown (Fig. 7F, G), this represents an appealing—but as yet unproven—pathway linking HNF4α to ER homeostasis. A major function of mature hepatocytes is the regulation of whole-body metabolism, for which TCA cycle activity is essential (50). In addition to their own metabolic needs, hepatocytes supply glucose to other organs during fasting, and that glucose is derived by the carboxylation of pyruvate to oxaloacetate and subsequent cataplerosis of oxaloacetate. Likewise, hepatic lipogenesis during nutrient abundance arises from cataplerosis of citrate. Pyruvate Dehydrogenase exerts a strong influence on TCA cycle activity by controlling the availability of Acetyl-CoA for condensation with oxaloacetate, and inhibition of PDK stimulates TCA cycle flux (51, 52). In our previous work, we showed that production of NADPH by the TCA cycle favors glutathione reduction, leaving the ER relatively hypo-oxidized (16). Since ER homeostasis is known to be exquisitely sensitive to reducing conditions in the organelle (53), upregulation of PDK4 (along with PDK2, the most highly expressed hepatic PDK isozymes (54)) could thereby alleviate ER stress through the suppression of TCA cycle activity. We found that NADPH was elevated in 5C-treated cells, in support of this model (data not shown), and are in the process of investigating this possibility more thoroughly. The relationship shown here between HNF4α and PDK4 is in contrast to the stimulatory effect of HNF4α overexpression on a *Pdk4* reporter in HepG2 cells (55). The reason for this discrepancy is unclear but could be accounted for if the effects of HNF4α on *Pdk4* expression depend on nutritional state or other conditions. Given that hepatocytes maintain metabolic flexibility to adapt to the nutritional needs of the organism, it seems logical that HNF4α, the master regulator of hepatocyte specification, would control *Pdk4* expression at some level.

A key question is how permanent the apparent interruption of hepatocyte identity is in hepatocytes exposed to chronic ER stress, and whether that phenotype underlies at least a portion of the liver injury seen in conditions like obesity, alcoholism, exposure to toxins, etc. that have been shown to have an ER stress component (3). In this study, we have referred to the gene expression changes that accompany ER stress in the liver as a loss of hepatocyte identity rather than as dedifferentiation, simply because we found no evidence that ER stressed hepatocytes achieve a truly mesenchymal fate nor that they transdifferentiate into, for example, cholangiocytes. The transience of a single tunicamycin challenge in a wild-type animal might be insufficient to truly effect a permanent cell fate change. However, it is worth noting that Cluster 7 genes in particular—those that most strongly encode hepatocyte identity and are most significantly tied to HNF4α—are also the most profoundly suppressed at a longer timepoint when the adaptive capacity of the UPR is impaired by deletion of ATF6α (Figure 1C). These genes are not ATF6α targets *per se*, in which case their regulation by ER stress would be lost when ATF6α is deleted. Rather, their enhanced suppression in *Atf6α*−/− animals is most likely a consequence of the fact that, without full upregulation of ER chaperones and other protective factors, *Atf6α*−/− cells experience much more persistent ER stress in the face of a particular challenge (28). This finding ties hepatocyte identity proportionally to the persistence of ER stress being experienced by the cell. It suggests that ER stresses that are truly chronic in nature might result in the permanent dedifferentiation of at least a subset of hepatocytes. The consequence of this dedifferentiation would be at best an impairment of normal liver function and at worst an increased propensity toward hepatocellular transformation. The association of dedifferentiation with exacerbated ER stress has also been observed in other cell types, most notably secretory cells including pancreatic beta cells (56–58) and thyrocytes (59), suggesting that it could be a conserved feature of UPR signaling in professional secretory cells.

It is widely accepted that ER stress contributes to liver injury through hepatocyte cell death, which causes inflammatory signaling that ultimately stimulates liver fibrosis and transformation (60). Indeed, during persistent stress cells will presumably be more likely to become overwhelmed by the stress burden and to die. And acute exposure to particularly hepatotoxic substances such as acetaminophen might result in a substantial level of ER stress-dependent cell death (61). However, for the most common forms of chronic liver injury including obesity and alcoholism, it is conceivable that the transcriptional and presumptively adaptive response to ER stress might contribute as much or more to liver dysfunction as does cell death. Understanding if UPR-dependent hepatocyte dedifferentiation contributes to such liver diseases will require a model of chronic ER stress that isolates the ER stress component from other confounding influences in a way that stressors such as a western diet or ethanol feeding do not.

### Limitations of the Study

Our work has demonstrated a bidirectional relationship between ER stress and HNF4α: ER stress disrupts the HNF4α-dependent GRN, and loss of HNF4α desensitizes hepatocytes to ER stress, at least caused by tunicamycin. Complementary model systems point to several possible pathways by which loss of HNF4α reduces the sensitivity of hepatocytes to ER stress, but we cannot yet definitively confirm or exclude any of these, likely in part due to the limitations of primary hepatocytes *in vitro*. We were also unable to successfully overexpress HNF4α *in vivo*—possibly due to its already very high expression—so we do not yet know whether HNF4α overexpression is sufficient to block the changes in expression of hepatocyte identity genes caused by ER stress. In addition, we cannot yet be certain whether the differential stress sensitivity seen in the *in vivo* deletion of HNF4α and in *in vitro* manipulation of hepatocyte identity through 5C treatment reflect the effects of the same cellular processes, or merely give concordant results coincidentally. We also do not know whether the differences in stress sensitivity between null- and 5C-treated cells are due to the latter remaining in a more differentiated state, or instead reflect specific effects of one or more of the 5C compounds on ER homeostasis independent of the cells’ differentiation state. Further investigation is required to address these issues.

## Materials and Methods

### k-means clustering

Microarray data were compiled from separate published experiments in which matched wild-type or *Atf6α−/−* (10), *Ire1α^LKO^* (13), or *Perk^LKO^* (11) animals were challenged with 1 mg/kg TM or vehicle for 8h, or in which *Atf6α−/−* or wild-type animals were challenged with 1 mg/kg TM for 34h (10). For each probe set, raw expression values were log_2_ transformed and were normalized against the average expression level for vehicle-treated wild-type animals. The data were then subjected to k-means clustering using the SimpleKMeans algorithm built into the Waikato Environment for Knowledge Analysis (Weka) machine learning software workbench (62). The distances between clusters and instances were computed using Euclidean distance. The genes in each cluster were subjected to pathway analysis using DAVID (63) and mined for enriched transcription factor binding sites using oPOSSUM (64). Some genes were represented by more than one probe set, and occasionally those probe sets showed discrepant behavior.

### Animal experiments

All animal procedures were approved by the University of Iowa Institutional Animal Care and Use Committee (IACUC). *Hnf4α^fl/fl^* (33) mice were housed in a pathogen-free facility at the University of Iowa on a 12-h light/ dark cycle. AAV8 virus was suspended in phosphate-buffered saline and injected intraperitoneally. TM or vehicle (DMSO) was dissolved in phosphate-buffered saline and was injected intraperitoneally. After 14 or 48h, liver tissues were harvested at the indicated time points after injections and either flash frozen or fixed in formalin (for histological analysis) and glutaraldehyde (for TEM). *Hnf4α*^fl/fl^ mice were injected with AAV8-TBG-iCre (Vector Biolabs, cat# VB1724) or AAV8-TBG-eGFP (Vector Biolabs, cat# VB1743) intraperitoneally (IP) using either 2.5 x10^11^ GC/mice or 5 x 10^11^ GC/mice. One week after virus injection, mice were fasted for 4 hours before IP injection with 0.5 mg/kg tunicamycin (TM) diluted in PBS or an equivalent amount of DMSO diluted in PBS. 14 hours after TM injection, the liver was harvested and either flash frozen for isolation of protein or RNA or fixed for histology or TEM analysis. Blood was collected via the retro-orbital route. AST (Biovision K753) and ALT (Biovision K752) values were determined using the manufacturer’s protocol. Protein was isolated by homogenization of samples in RIPA buffer, and RNA by homogenization in Trizol (ThermoFisher), and qRT-PCR (including piloting of primer sets to confirm efficiency and specificity), immunoblot, and *Xbp1* splicing assays were as described (16, 65). qRT-PCR C_t_ values were normalized against the average of two housekeeping genes (*Btf3* and *Ppia*) to mitigate the possibility of changes in a housekeeping gene confounding the expression quantification. The following antibodies were used for blotting, which controls for specificity as indicated: HNF4α (Abcam ab181604 or ab201460; knockout), BiP (BD Biosciences 610978; overexpression), CHOP (ProteinTech 15204-1-AP; knockout), Calnexin (Assay Designs ADI-SPA-865-F; migration upon protease digestion of microsomes), ADRP (NB110-40877; immunostaining of lipid droplets), β-actin (Cell Signaling 4970) TRAPα (provided by R.S. Hegde; recognition of purified protein), anti-mouse HRP (Jackson Immune Research, 115-035-003) and anti-rabbit HRP (Jackson Immune Research, 111-035-045). Ki67 staining was performed by the University of Iowa Comparative Pathology Laboratory.

### RNA Seq Analysis

RNA-seq was performed by the Genomics Division, Iowa Institute of Human Genetics (IIHG) Core, University of Iowa. Following isolation, RNA was further purified by using an RNeasy Mini Kit (Qiagen), quantified using a Qubit machine (ThermoFisher), and 2 μg per sample was used for sequencing on an Illumina HiSeq 4000/75PE Sequencer, generating 23 million reads per sample. RNA seq data quality was analyzed using FastQC (https://www.bioinformatics.babraham.ac.uk/projects/fastqc/, v.0.11.8). The FASTQ RNA-seq files were aligned to Ensembles Transcriptomes v.94. for *Mus musculus* (https://useast.ensembl.org/info/data/ftp/index.html). A pseudo alignment program *Kallisto* [v_0.45.0; (66)] was used to align the RNA seq FastQC data files to determine the transcripts abundances. Downstream differential gene expression analysis was performed by *DESeq2* [v_1.34.0 (67) using the *R Studio* statistical package (v_4.1.2). All the dependencies required for the analysis of RNA seq data were gathered from the Bioconductor site (https://www.bioconductor.org). Gene set enrichment was performed by using GSEA software (http://www.gsea-msigdb.org/gsea/index.jsp; GSEA_4.0.3) following the pre-ranked method. Normalized differentially expressed genes from DESeq2 analysis were ranked based on their log_2_FC. The hallmark gene v7.1 for pre-ranked method and gene symbol remapping 7.1. and a default setting was used for analysis.

### Primary hepatocyte culture

Primary hepatocytes were isolated and cultured as described (16). Long-term primary hepatocytes culture maintenance media was prepared following (37). Null media was prepared by supplementing William’s E media with B27 (50X; Gibco,17504044), Pen. /Strep (Gibco, 15140-122), Amphotericin B (Thermo, 15290-018), and Gluta-max (Gibco, 35050-061). Dedifferentiation inhibition media (5C media) was prepared by supplementing null media with Forskolin (20uM; Enzo, BML-CN100), SB431542 (10uM; Tocris, 1614), IWP2 (0.5uM; Tocris, 3533), DAPT (5uM; Tocris, 2634), and LDN193189 (0.1uM; Tocris, 6053). After isolation as above, cells were resuspended in null media and plated on collagen-coated plates. Four hours after isolation, fresh null media was replaced. On the next day and every 24 hours thereafter, Null or 5C media was added. In our experience, the source of collagen used to coat the plates (C3867, Sigma) was an essential feature. Cultured primary hepatocytes were treated with ER stressors – tunicamycin (5 μg/ml), thapsigargin (50 nM), or vehicle (DMSO) for 16hrs. For knockdowns, dsiRNAs (IDT, Coralville, IA) or non-targeting controls were used. Cells were transfected using Dharmafect-4 reagent (T-2004-03) according to the manufacturer’s instructions. Cells were incubated in transfection media for 36-48hrs and 1mL of fresh primary hepatocyte complete media was added as necessary. Sequences of *Pdk2* and *Pdk4* dsiRNAs were: *Pdk4*: AAAGAAGACCUUACAAUCAAGAUTT, GGAAUCAAAGCACUUUAUCFAGCTT; *Pdk2*: AGAUCAGAACUGUCUGUUUUCUACT, AGUAUUACAUGGCUUCCCCUUGACCT. For both genes, both dsiRNA were independently validated for specificity and produced equivalent results.

### Histological and analysis and electron microscopy

Immunohistochemistry (IHC) of samples fixed in 4% neutral buffer formalin-fixed liver tissue was performed by the Comparative Pathology Laboratory (CPL), University of Iowa. For TEM, a small piece of liver tissue was finely minced and fixed using 2.5% glutaraldehyde (provided by the microscopy core) and stored at 4°C. Samples were further processed for TEM images by the Central Microscopy Core, University of Iowa. Analysis of the images was performed double-blinded. Samples were blinded when submitted to the core and further images obtained from the core were again blinded. Image J software (v.1.8.0_172) was used to quantify the circularity of the organelle. A value close of 1 represents a circular organelle, and a value close to zero (0) is considered non-circular.

### Statistics

Prism software (v.9.) was used for the statistical analysis. Except where noted otherwise, one-way ANOVA was used with *Tukey’s* post hoc correction for performing multiple comparisons. “n” numbers represent either individual mice (for *in vivo* experiments) or independently treated wells (for cell culture experiments).

### qRT-PCR sequences

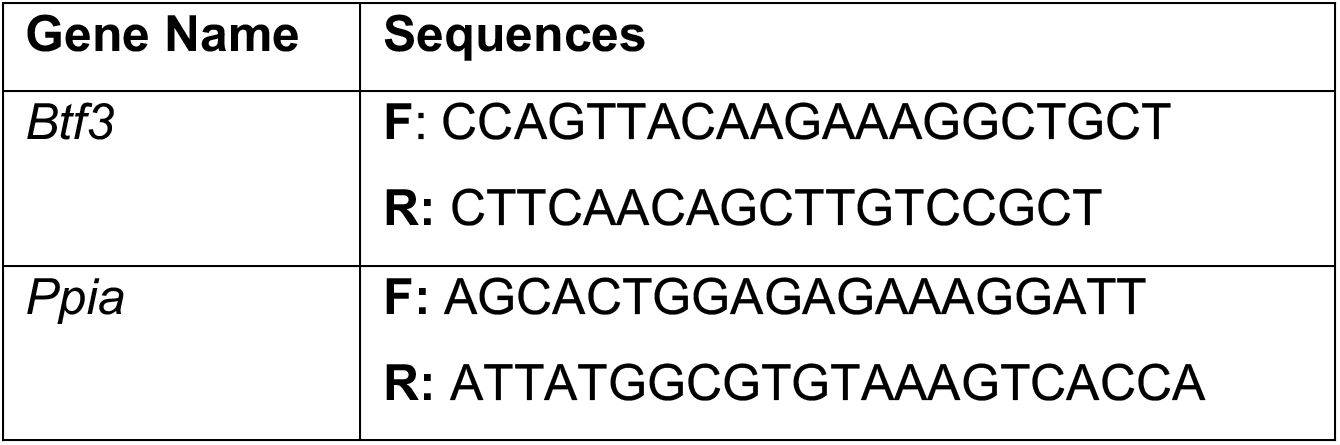
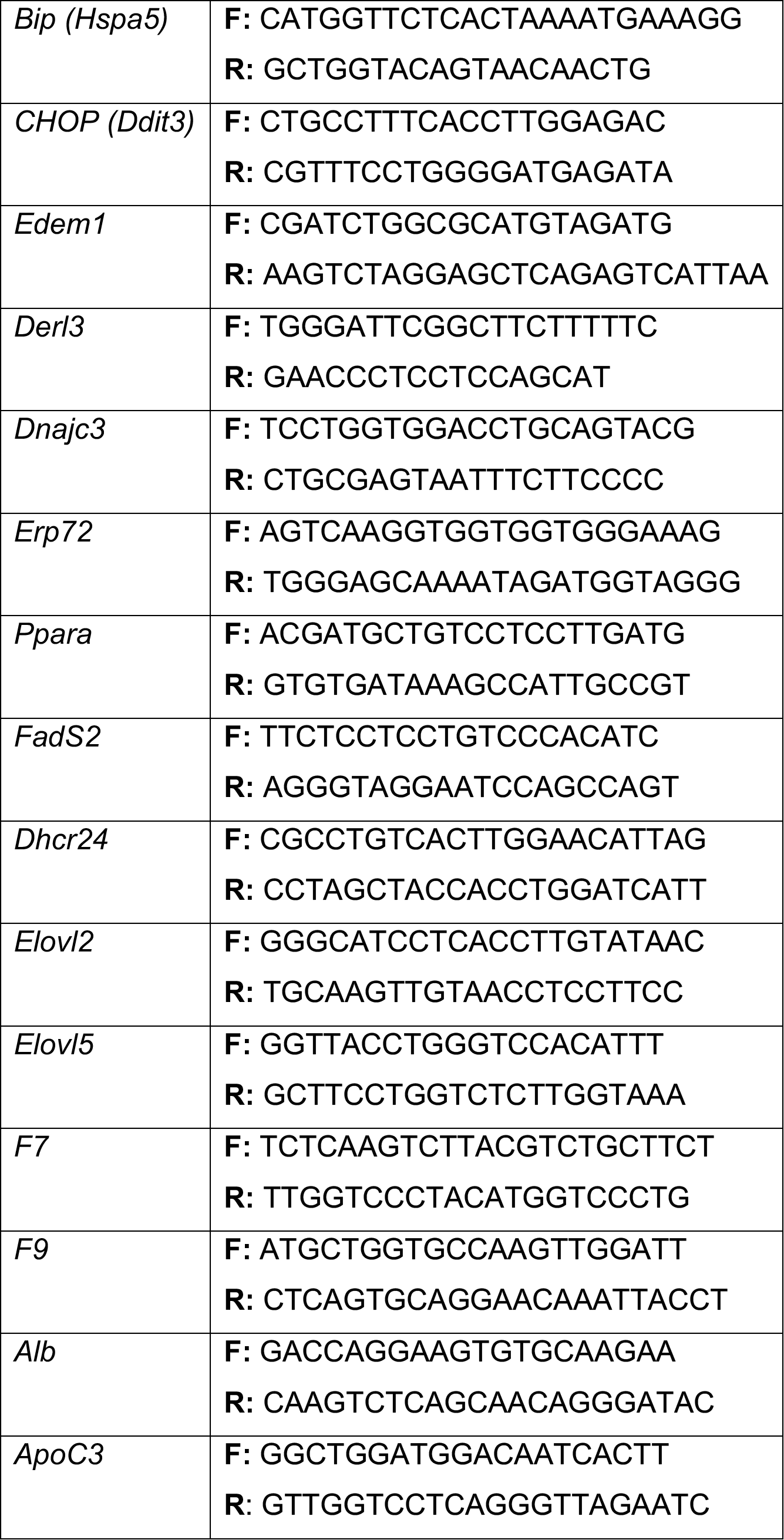
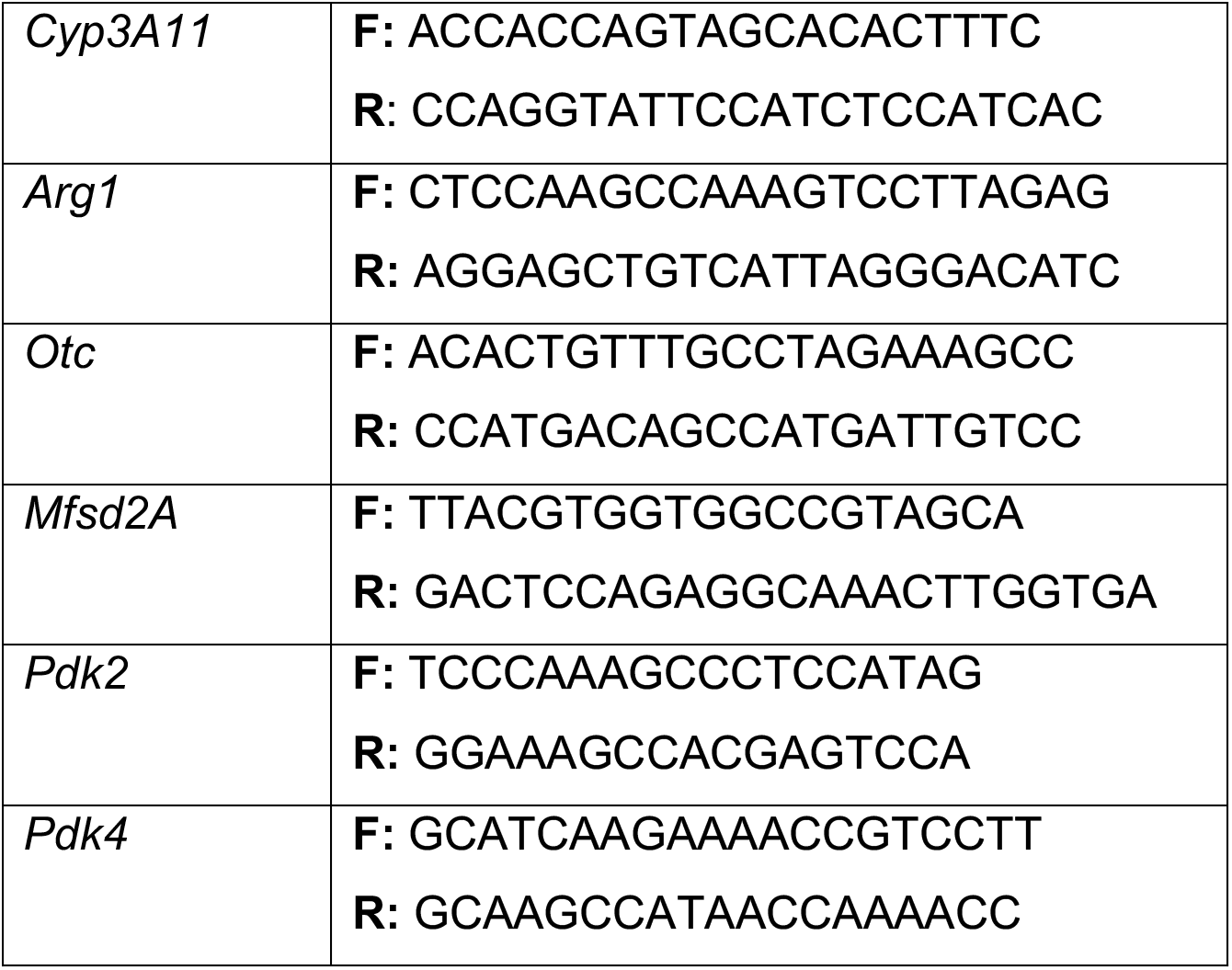

## Acknowledgments

We thank Ling Yang, Qingwen Chen, and Mark Li, University of Iowa, for technical assistance and Holger Willenbring, University of California San Francisco, for enlightening discussions. We also thank Ling Yang, Matthew Potthoff, Vitor Lira, and Robert Cornell, University of Iowa, for fruitful discussions. This work was funded by grants R01 GM115424 to DTR, F31 DK130250 to ERG, and R01 DK98414 and R56 DK112768 to UA.

## Author Contributions

AS and DTR conceived and designed the experiments. AS performed the majority of the experiments. IH performed the *in vivo* experiment from which RNA-seq data were generated, with UA. KD provided guidance on RNA-seq analysis to AS. MS performed *k-means* clustering. AS and DTR wrote the manuscript. All authors read and approved the manuscript.

