## Supplementary Figures for "Interference with the HNF4-dependent gene regulatory network diminishes ER stress in hepatocytes"

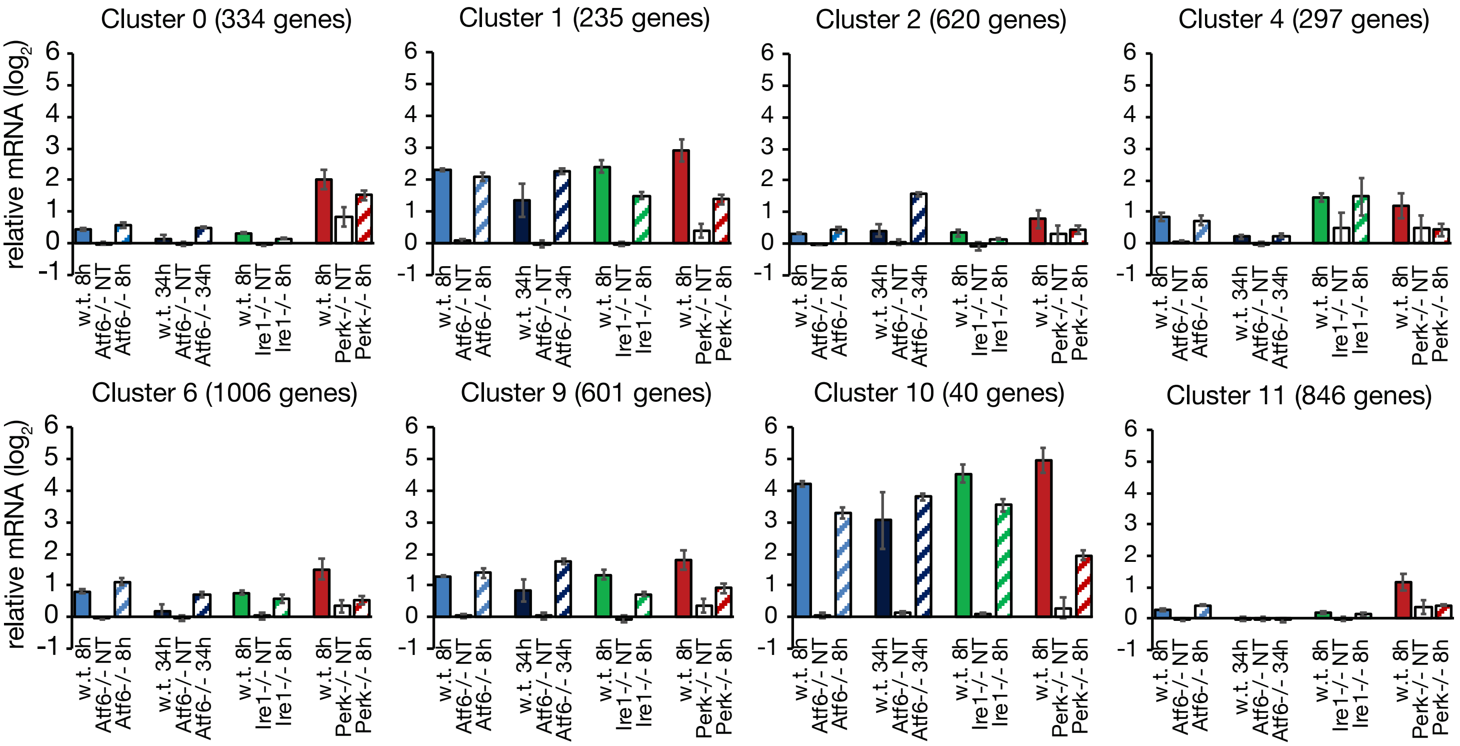


**Figure S1. Centroids for upregulated genes from k-means clustering**

Genes that were upregulated by ER stress from the microarray datasets were partitioned into 8 of the 12 clusters. The centroids for these genes and the number of genes in each cluster are shown here.

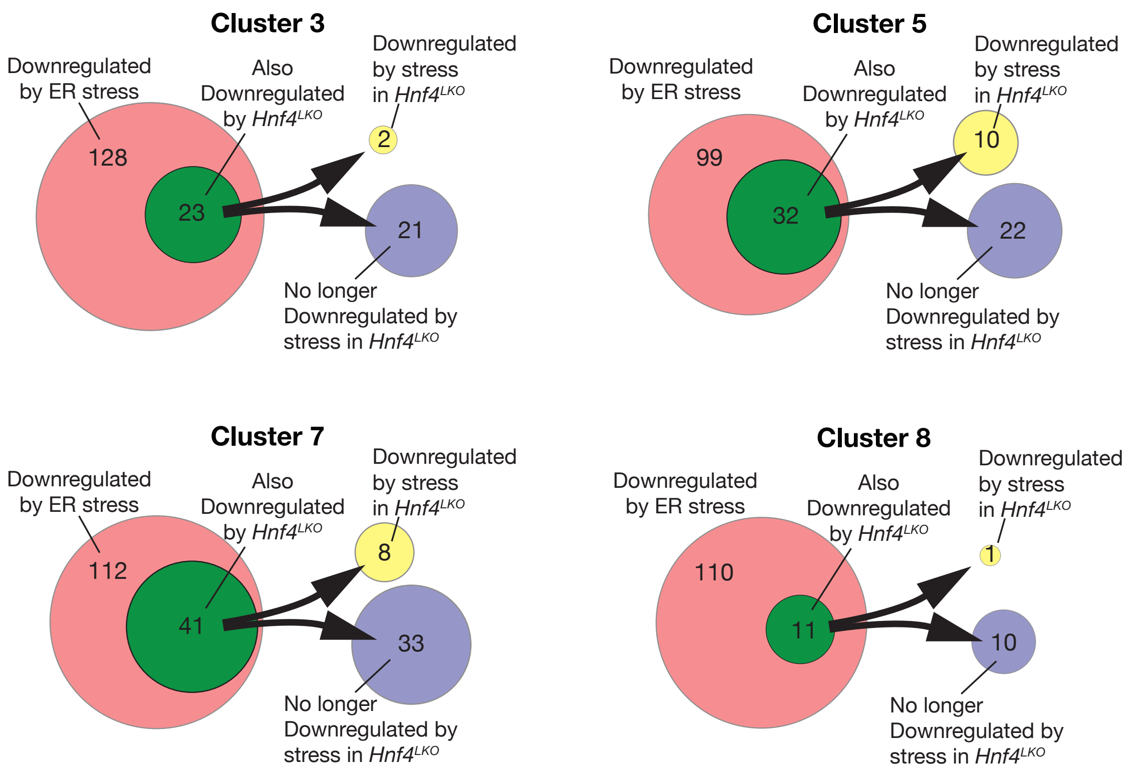


**Figure S2. Stress regulation of HNF4α-dependent genes from clusters 3, 5, 7, and 8**

Red circles represent genes from each cluster that were confirmed to be significantly downregulated by TM treatment in wild-type animals from RNA-seq dataset. Green circles represent the genes within that group that were also significantly downregulated by loss of HNF4α. The genes of that group were subdivided into those that were no longer downregualted by TM in $Hnf4\alpha^{LKO}$ animals (blue circles) and those that were further downregulated in the same conditions (yellow circles).

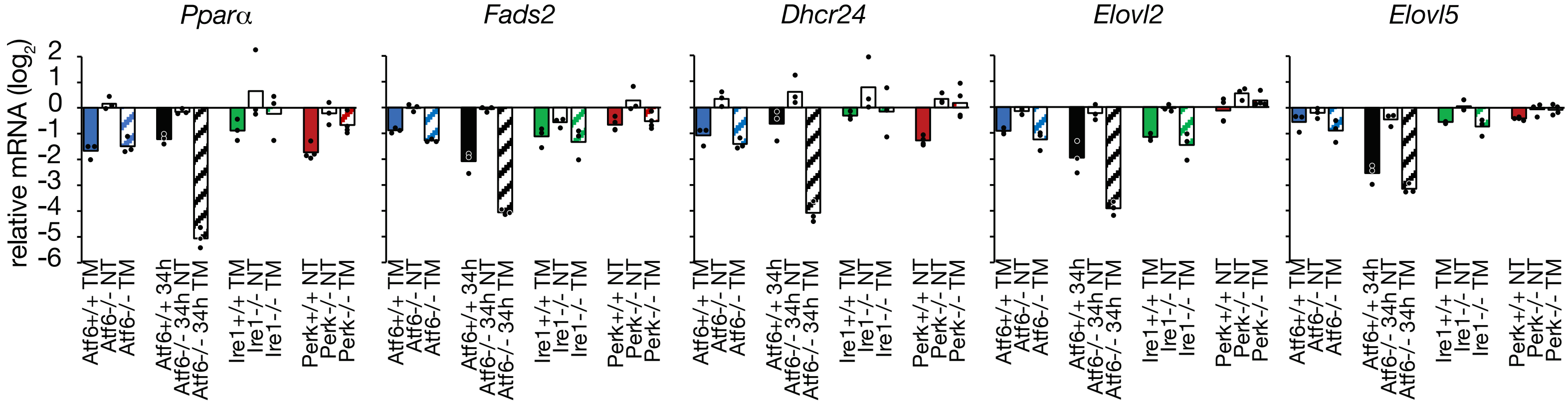


**Figure S3. Expression of selected cluster 7 genes in microarray data**

The behavior of the 5 cluster 7 genes shown in Figure 3B in the microarray datasets from Figure 1 is shown here.

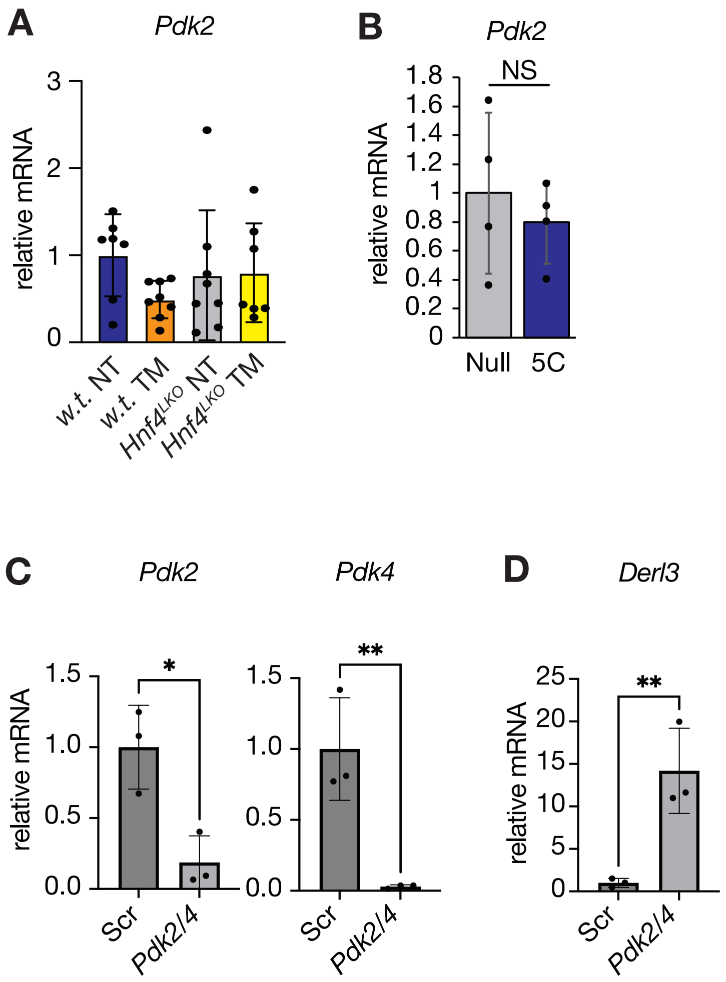


**Figure S4. Pdk2/4 knockdown**

**(A, B)** Expression of *Pdk2* determined by qRT-PCR in vivo (A) and in vitro (B).
**(C)** qRT-PCR shows effective knockdown of *Pdk2* and *Pdk4* by siRNA
**(D)** Similarly to *Bip* and *Chop*, *Derl3* mRNA is upregulated by *Pdk2/4* knockdown.
